# Normative Growth Modeling of Cortical Thickness Identify Neuroanatomical Variability and Distinct Subtypes in Brainstem Tumor Patients

**DOI:** 10.1101/2024.08.01.606270

**Authors:** Heyuan Jia, Kaikai Wang, Peng Zhang, Mingxin Zhang, Yiying Mai, Congying Chu, Xuntao Yin, Lingzhong Fan, Liwei Zhang

**Affiliations:** School of Instrumentation and Optoelectronic Engineering, Beihang University, Beijing, China; Institute of Large-scale Scientific Facility and Centre for Zero Magnetic Field Science, Beihang University; Brainnetome Center, Institute of Automation, Chinese Academy of Sciences, Beijing 100190, China; Department of Neurosurgery, Beijing Tiantan Hospital, Capital Medical University, Beijing, China; Department of Radiology, Guangzhou Women and Children’s Medical Center, Guangzhou Medical University, Guangzhou, China; School of Health and Life Sciences, University of Health and Rehabilitation Sciences, Qingdao 266000, China; China National Clinical Research Center for Neurological Diseases, Beijing Tiantan Hospital, Capital Medical University, Beijing, China; Beijing Neurosurgical Institute, Beijing Tiantan Hospital, Capital Medical University, Beijing 100070, China

**Keywords:** brainstem tumors, cortical thickness, normative model, subtype classification, tumor growth pattern

## Abstract

**Background:** Brainstem tumors can cause structural brain changes, but the resulting heterogeneity within wholebrain structure is not well-studied. This study examines cortical thickness to identify patterns of structural alterations and explore underlying biological subtypes and their associations with clinical factors.

**Materials and Methods:** This study involved 124 pediatric brainstem tumor patients, aged 4-18 years. Cortical thickness was measured using CAT12 segmentation of 3D T1-weighted structural MRI. A normative model was established using data from 849 healthy children. Deviations in cortical thickness were estimated, and patients were classified into two subtypes using spectral clustering. Clinical statistical analyses were conducted with SPSS 26.0.

**Results:** The normative model revealed significant heterogeneity in cortical thickness deviations, which correlated with tumor size and growth patterns. Focal tumors primarily caused negative deviations (t = 3.14, p = 0.02). There was a significant positive correlation between extreme positive deviations and tumor volume (r = 0.214, p = 0.010), and between extreme negative deviations and progression-free survival (r = 0.39, p = 0.008). Two subtypes were identified: Subtype 1, consisting of diffuse tumors with extreme positive deviations, and Subtype 2, consisting of focal tumors with extreme negative deviations. Subtype and tumor growth pattern significantly influenced duration (p < 0.01). The Kaplan-Meier survival curves for Subtype 1 and Subtype 2 demonstrated a significant difference in survival probabilities over time (p = 0.03).

**Conclusion:** Overall, this study identifies two major patterns of cortical thickness changes in brainstem tumor patients, enhancing our understanding of their relationship with cortical morphology. The findings suggest that cortical thickness alterations could serve as valuable biomarkers for predicting progression-free survival, which is crucial for clinical assessment and personalized treatment strategies. This research provides new insights into the physiological mechanisms by which brainstem tumors affect brain structure, supporting more precise clinical interventions and efficacy monitoring in the future.

## Introdution

Brainstem tumors, though relatively rare, are prevalent among pediatric brain tumors and typically arise in the midbrain, pons, and medulla. These tumors include various pathological types, such as astrocytomas, gangliogliomas, ependymomas, and oligodendrogliomas, and are categorized into diffuse and focal types based on imaging characteristics(Lee et al, 1985; Recinos et al, 2007). Diffuse brainstem tumors are highly invasive and fast-growing, leading to rapid symptom onset like cranial nerve deficits, ataxia, and long tract signs, resulting in a poor prognosis due to the difficulty of surgical removal(Chou et al, 2024). Conversely, focal brainstem tumors grow more slowly, are well-defined, and primarily cause symptoms through local compression. They are more amenable to surgical resection, offering a better prognosis(Klimo et al, 2013). Usually, diffuse tumors are aggressive and progress quickly, while focal tumors are focal and develop symptoms gradually.

The heterogeneity of brainstem tumors is evident not only in their pathology and clinical presentation but also in their treatment response. Current research mainly focuses on the tumor and its surrounding tissues locally(Boukaka et al, 2023). However, studies on the global changes induced by these tumors are relatively scarce. Given the brainstem’s critical role as a conduit between the brain and the spinal cord, tumors in this region are likely to disrupt essential anatomical pathways(Smith & DeMyer, 2003). This disruption can lead to corticospinal disconnection, where communication between the cortex and the spinal cord is impaired. Such disconnection may result in widespread cortical atrophy, as the absence of proper signaling and feedback mechanisms can lead to degeneration in cortical regions dependent on these pathways for normal function(Acharya et al, 2022). Brainstem injuries can affect other significant pathways within the central nervous system. They can interrupt sensory and motor pathways, leading to deficits in sensation and movement(Newman, 1995). Disturbances in autonomic pathways can impact vital functions like respiration and cardiac regulation(Novak et al, 1995). These broad effects highlight the need for comprehensive studies. Such studies should address both the local tumor environment and the wider implications for brain function and structure.

Brain tumors, particularly gliomas, are seen as systemic diseases. Glioma cells infiltrate the brain along white matter tracts, affecting cortical structures(Duffau, 2014; Wang et al, 2022). Midline diffuse gliomas cause cortical thickening and demyelination, indicating brainstem tumors can induce widespread changes. These changes impact cognitive functions and recovery post-surgery(Zhang et al, 2024). Cortical alterations correlate with cognitive impairments like executive function and memory deficits(Osswald et al, 2015). Understanding brainstem tumors’ effects on distant brain regions is crucial for comprehensive treatment(Chieffo et al, 2023).

Neuroplasticity research indicates that the brain undergoes cortical remodeling in response to injury and stress. This remodeling results in compensatory and decompensatory patterns(Wittenberg, 2010). For example, corticospinal disconnection might cause cortical atrophy in multiple brain regions, representing a form of decompensation(Pistoia et al, 2016). Conversely, certain brain areas may compensate for the loss of function by increasing neuronal connections or altering function(Osswald et al, 2015). Given their unique growth location and pattern, brainstem tumors likely induce both types of cortical changes. We hypothesize that brainstem tumor patients exhibit significant heterogeneity in cortical thickness changes. This heterogeneity is closely related to the tumor’s growth location and pattern. Different growth locations and patterns may lead to distinct cortical thickness alteration patterns. These changes can subsequently affect neurological function and cognitive performance. This hypothesis offers a new perspective for studying the global impact of brainstem tumors on the brain.

To address this hypothesis, we collected multisite, high-quality strcutural MRI data from a large sample of 849 children aged 4-18 years. An overview of the study workflow is shown in Figure 1. We first established normative growth model of cortical thickness in 210 brain regions to characterize the neuroanatomical heterogeneity of patients with brainstem tumor. The cortical thickness signature was used for normative modeling since it reflects the neural cytoarchitecture and is a valuabel statistical approach.

**Fig 1.**
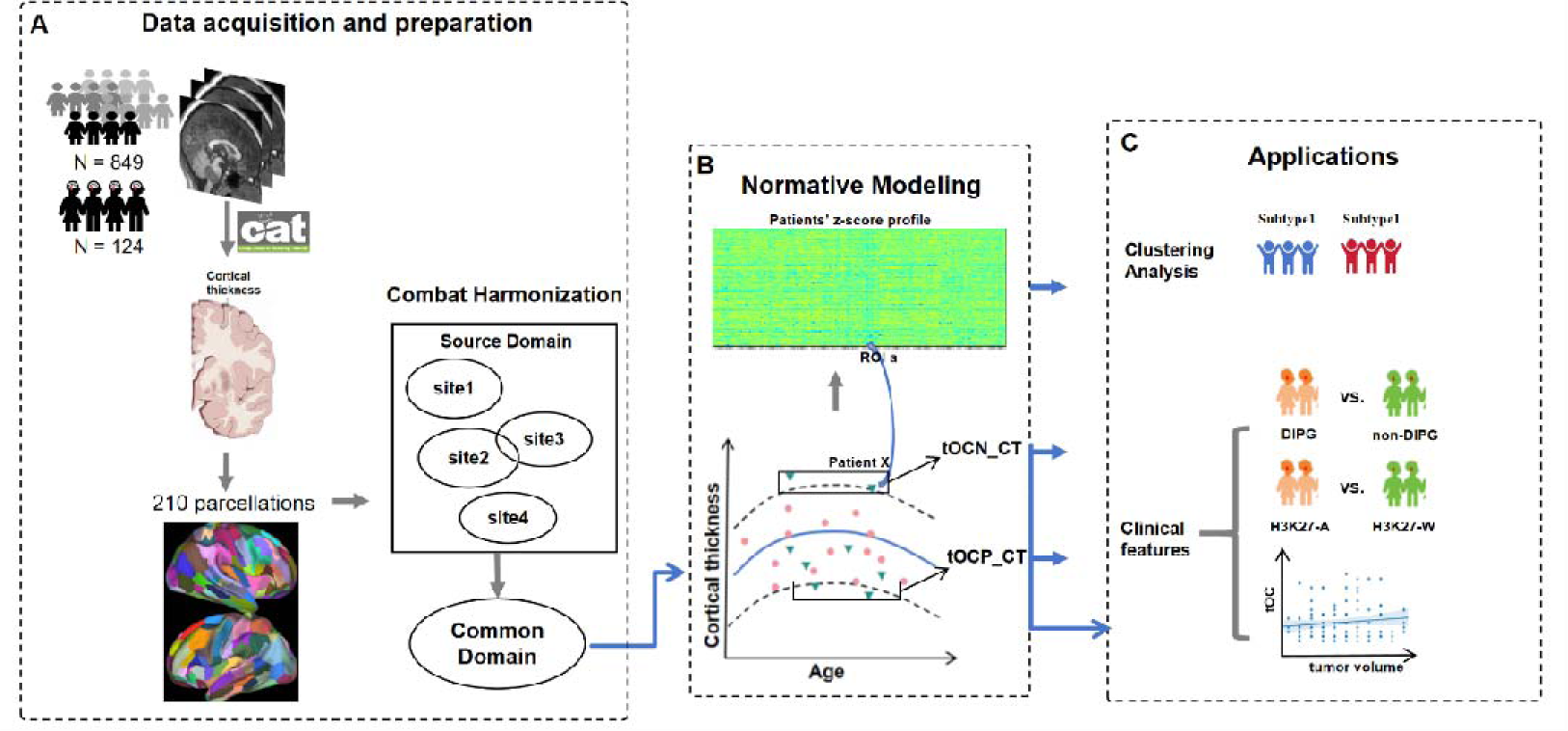
Flow chart of cortical thickness heterogeneity analysis and subtype identification. (A) Acquisition and preprocessing of structural MRI (sMRI) data, including cortical thickness segmentation, volume extraction of Regions of Interest (ROI), and data harmonization. (B) Establishment of a normative model using the Gaussian Process Regression (GPR) method, followed by the calculation of individualized cortical thickness (CT) deviations for patients. (C) Application of individualized CT deviation data to identify subtypes using spectral clustering, and the subsequent analysis of clinical differences among these subtypes.

To validate our hypothesis, we meticulously designed the research process illustrated in Figure 1. We gathered high-resolution structural MRI data from multiple centers, involving 849 healthy children aged 4 to 18 years. Using this data, we constructed a normative model based on cortical thickness measurements from 210 brain regions, providing a crucial reference for assessing neuroanatomical heterogeneity in brainstem tumor patients. Our model allows precise assessment of individualized deviations in cortical thickness. We employed spectral clustering to identify subtypes of tumor patients and analyzed differences in tumor growth patterns, disease progression, and tumor locations among these subtypes. This approach enhances understanding of the diverse impacts of brainstem tumors and aids in tailoring patient-specific treatment strategies.

## Material and methods

### Participants

A cohort of 124 pediatric patients with brainstem tumors was studied at Beijing Tiantan Hospital from April 2020 to December 2023. Criteria for inclusion were: new diagnosis without prior treatment, tumors confined to the brainstem, no psycho-neurological or behavioral disorders, and no developmental disorders. All participants underwent 3DT1 MRI, with informed consent obtained from parents under an IRB-approved protocol.

To create a normative model, data from 849 healthy children were sourced from four databases: 61 local recruits in Beijing, 69 from the fcon_1000 projects dataset, 450 from the Child Connectome Project (devCCNP)(Xi-Nian & Consortium, 2024), and 269 from the Children’s Hospital in Guangzhou. MRI T1 images and phenotypic data were downloaded and summarized. After excluding those with severe cerebellar compression or poor image quality, the final analysis included 124 patients and 849 healthy children.

### MRI acquisition and processing

All participants underwent a structural MRI using a 3.0 T Ingenia CX scanner (Philips Healthcare, Best, Netherlands) with a 32-channel head coil. The scan included a whole-brain 3D T1-weighted anatomical sequence with 196 contiguous sagittal slices, each with a voxel size of 1.0 x 1.0 x 1.0 mm³. The parameters for the 3D-T1 sequence were: repetition time (TR) =6.572ms; echo time (TE) =3.025 ms; flip angle (FA) = 8°; slice thickness = 1mm; in-plane resolution = 1*1mm.

For each individual, cortical thickness was calculated using the CAT12 toolbox (version 12.7). The procedure involved several steps: MRI data were first segmented to distinguish between gray matter, white matter, and cerebrospinal fluid. Following segmentation, the pial surface and the gray matter/white matter boundary surfaces were reconstructed. Cortical thickness was then measured as the distance between corresponding vertices on these surfaces(Keller & Roberts, 2008). The brain was parcellated into 210 distinct regions using the Human Brainnetome Atlas, and the mean cortical thickness for each region was calculated(Fan et al, 2016).

### Normative modeling

To harmonize MRI data across three datasets and address inter-scanner variability, we employed the Combat method, which has proven effective in reducing non-biological variance from site differences while maintaining biological variability(Fortin et al, 2018). This approach was applied to ensure uniformity across datasets, accounting for age, gender, and total intracranial volume (TIV) as covariates, while excluding diagnostic labels.

For normative modeling, we processed cortical thickness for 210 regions using Python 3.9 and PCNToolkit 0.22. We applied Gaussian Process Regression (GPR) to create region-specific normative models, incorporating age, gender and TIV as covariates. This Bayesian technique, implemented in PCNToolkit, accounts for prediction uncertainty and provides probability distributions for predicted volumes(Marquand et al, 2016). We validated these models through 10-fold cross-validation on the healthy control dataset before finalizing the normative models. These models were then used to predict cortical thickness for each participant and assess deviations.

To assess deviations in cortical thickness for glioma patients, we used percentile charts derived from healthy controls. We quantified deviations by comparing observed cortical thickness with those predicted by the normative model. Each region’s deviation was computed using the formula:

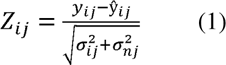

where y_ij_ is the cortical thickness, ŷ_ij_ is the expected cortical thickness estimated from the GPR, σ_ij_ is the predictive uncertainty, and σ_nj_ is the variance learned from the normative distribution n.

### Identifying Patients with Brainstem Biotypes using Spectral Clus tering

To identify patients with brainstem tumor biotypes, we performed spectral clustering analysis on 124 patients’ cortical deviation maps using the Scikit-learn. Spectral clustering was subsequently applied to the cortical deviation maps to cluster the participants into subgroups. The optimal number of clusters was determined according to Calinski-Harabaz Index(Bu et al, 2024).

### Outlier Definition and Calculation

Outliers indicating significant deviations in cortical thickness were identified using the absolute value of Z-scores. Regions with |*Z |* > 1.96 were classified as extreme deviations, representing the 2.5th percentiles at both ends of the normative distribution. For each brain region, the proportion of individuals with extreme positive deviations was defined as the extreme positive deviation rate, while the proportion of individuals with extreme negative deviations was defined as the extreme negative deviation rate. Each region’s deviation was computed using the formula

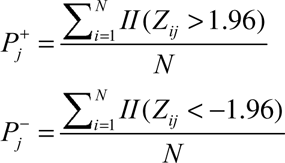

Specifically, let *N* be the total number of subjects, and let *II* (condition) be an indicator function that equals 1 if the condition is true and 0 otherwise.

These thresholds focus on identifying substantial deviations in cortical thickness. For each participant, we computed the total outlier count of negative deviation in cortical thickness (tOCN-CT) and the total outlier count of positive deviation in cortical thickness (tOCP-CT) across all 26 regions. This method allows for a comprehensive evaluation of both significant reductions and increases in cortical thickness associated with the condition.Each participant’s deviation was computed using the formula:

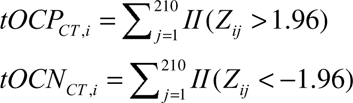

Specifically, *i* be the *i* -th subject, and *j* be the *j* -th brain region out of 210 brain regions, where *II* (condition) is an indicator function that equals 1 if the condition is true and 0 otherwise.

### Statistical Analysis

In our statistical analysis, we assessed the differences between two subtypes based on various factors by SPSS (version 26.0). For continuous variables such as total deviation rate of cortical thickness, tOCN-CT, tOCP-CT, clinical features like KPS (Karnofsky Performance Status), duration, tumor volume, and PFS (Progress Free Survival) we employed independent two-sample t-tests or Mann-Whitney U tests, depending on whether the data followed a normal distribution. For categorical variables, such as gender and tumor growth patterns, we used chi-square tests. To evaluate the interaction effects of tumor growth patterns and subtypes on clinical factors, we conducted two-way ANOVA. Statistical significance was determined with a two-tailed p-value of less than 0.05. Kaplan-Meier survival analysis was performed to estimate and compare the PFS between subtypes, and differences were assessed using the log-rank test.

## Results

### Cortical Thickness Heterogeneous Alterations in Partients with Brainstem Tumor

Utilizing normative growth models for cortical thickness across 210 brain regions, we investigated structural changes in patients with brainstem tumors(for model performance, see Table 1).

**Table 1.** GPR model performance indicator.

|  | SMSE | Rho |
| --- | --- | --- |
| Min | 0.11 | 0.10 |
| Max | 0.99 | 0.95 |
| Mean | 0.43 | 0.81 |

Our findings revealed an age-related thinning of cortical thickness in most regions, regardless of gender(Fig 2A, S1). Over 25% of patients showed extreme deviations in at least five brain regions. As the number of deviated regions surpassed ten, the proportion of patients with positive extreme deviations (increased cortical thickness) markedly decreased. Conversely, 13% of patients consistently exhibited extreme negative deviations (reduced cortical thickness), a phenomenon absent in healthy children. Remarkably, even with more than 25 deviated regions, about 1% of patients continued to show significant deviations. (|Z| > 1.96)(Fig 2B). This underscores the pronounced heterogeneity in cortical thickness characteristics among brainstem tumor patients. Mapping these extreme deviations onto brain cortex images further illuminated concentrated cortical thickening in the prefrontal lobe, precentral gyrus, postcentral gyrus, parietal lobe, and temporal pole, and widespread cortical atrophy in the inferior temporal gyrus, orbitofrontal gyrus, and occipital lobe (Fig 2C). Comparative analysis highlighted statistically significant differences in these regions between patients and healthy children (Fig 2D).

**Fig 2.**
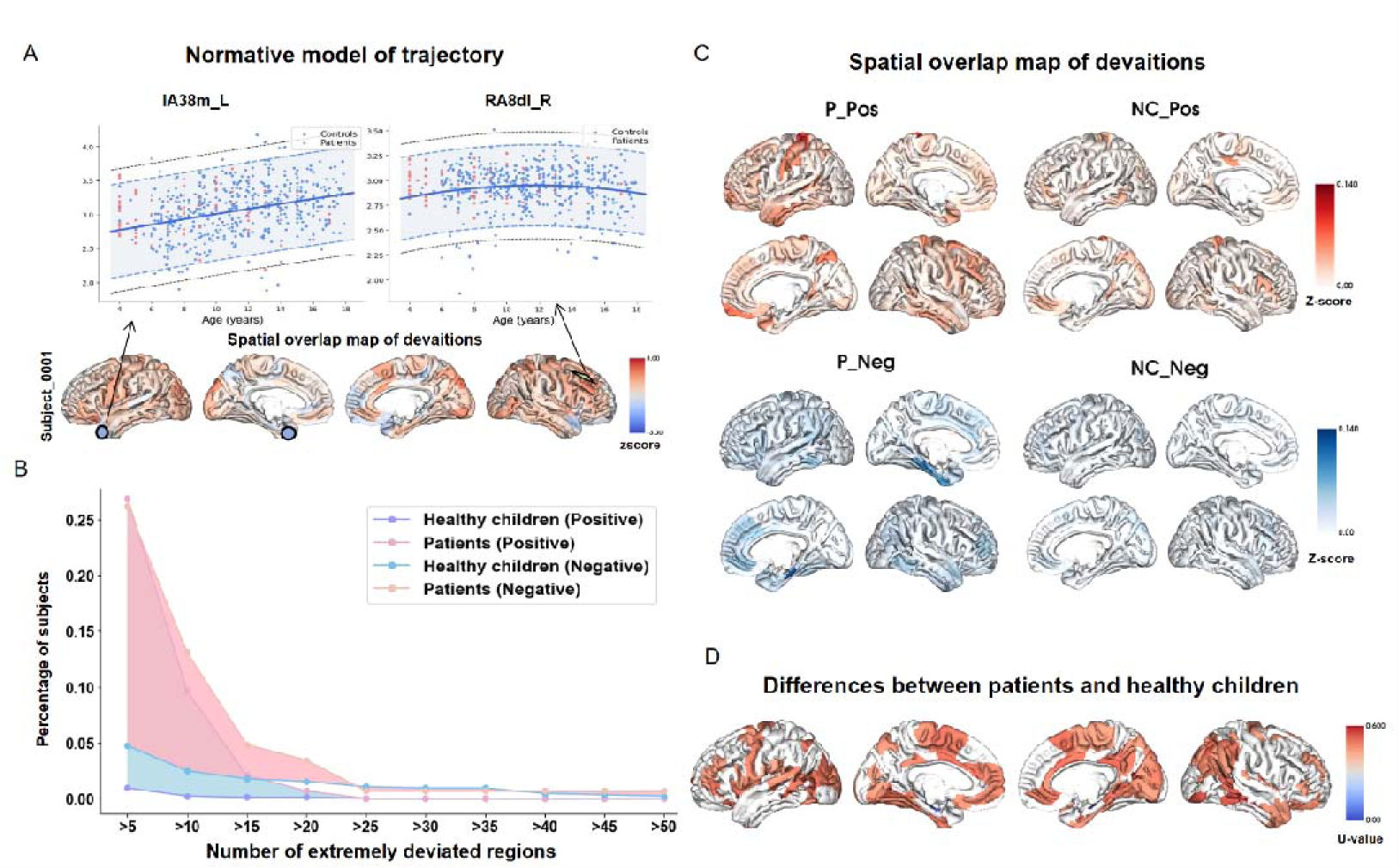
Heterogeneity Analysis of Cerebral Cortical Thickness in Patients with Brainstem Tumors. (A) Schematic diagram illustrating the curve fitting process for the Gaussian Process Regression (GPR) normative model. (B) Butterfly diagram depicting the extreme deviation rate across different brain regions. (C) Spatial overlap map showing areas of positive deviation (P_Pos) and negative deviation (P_Neg) in patients. (D) Comparative analysis highlighting differences in cortical thickness between patients and healthy children.

### Relationship between Cortical Thickness Variations and Clinical Characteristics in Patients with Brainstem Tumor

Through univariate analysis, we explored the relationship between individualized variations in cortical thickness and clinical characteristics in patients. Our results indicated that patients with focal tumors had significantly more negatively deviated brain regions compared to those with diffuse tumors (U = 1090.4, p = 0.04). However, no significant correlation was found between tumor growth pattern and the tOCP_CT (U = 1573, p = 0.44) (Fig 3A), suggesting that focal tumors are specifically associated with reduced cortical thickness. Additionally, Pearson correlation analysis revealed a positive correlation between tumor volume and tOCP_CT (r = 0.214, p = 0.010) (Fig 3B). Furthermore, Pearson correlation analysis demonstrated a positive correlation between tOCN-CT and PFS (r = 0.39, p = 0.008) (Fig 3C), indicating that higher tOCN-CT is associated with longer PFS. These findings underscore the influence of tumor size and growth pattern on cortical thickness variations in brainstem tumor patients.

**Fig 3.**
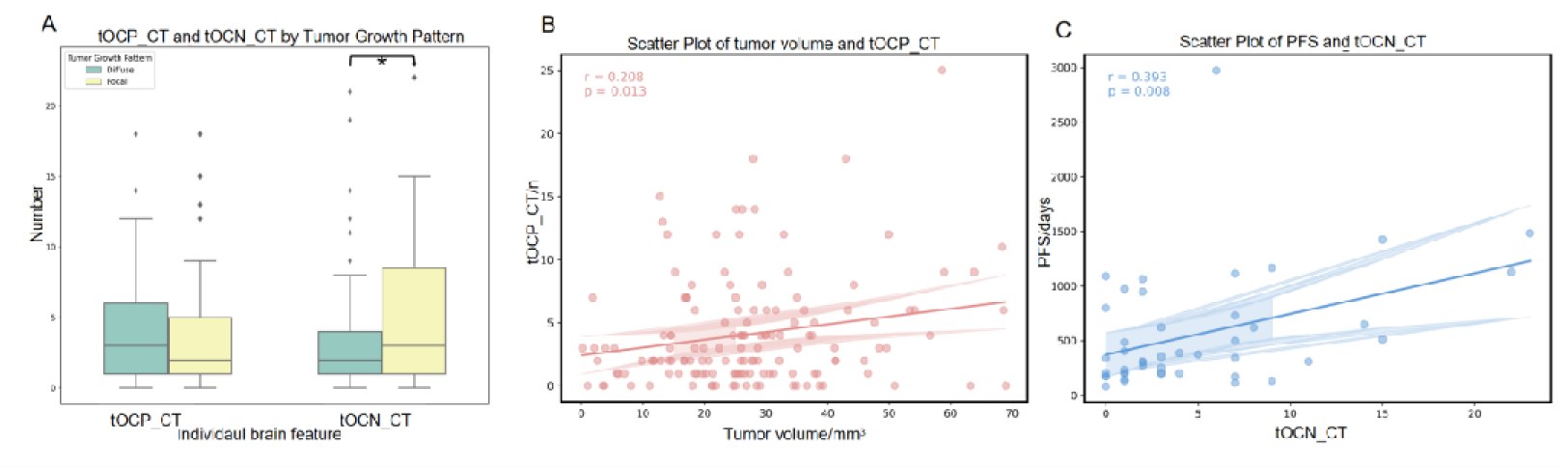
Clinical effects on cortical thickness deviation. (A) Differences in tOCP_CT and tOCN_CT between focal and diffuse tumors. (B) Correlation between tumor volume and tOCP_CT. (C) Correlation between PFS and tOCN_CT. PFS: progression-free survival, tOCN_CT: total outliers count of negative deviation of cortical thickness, tOCP_CT: total outliers count of positive deviation of cortical thickness. *: p<0.05.

### Identification of Two Subtypes in Patients with Brainstem Tumor

Employing spectral clustering analysis, we successfully classified brainstem tumor patients into two distinct biological subtypes based on their cortical thickness deviation patterns (see Table 2). 124 patients were divided into Subtype 1 (n = 93) and Subtype 2 (n = 31). The median age of the patients marked as Subtype 1 and Subtype 2 having median ages of 8.10 (SD = 3.44) and 9.39 (SD = 3.37) years. The gender distribution with Subtype 1 was 47 males and 46 females, and Subtype 2 was 17 males and 14 females.

**Table 2.** Characteristic differences between subtypes.

| Variations | Patients | Subtype 1 | Subtype 2 | p |
| --- | --- | --- | --- | --- |
| Age/years, median(std) | 8.19(3.39) | 7.47(3.17) | 8.5(3.46) | 0.14 |
| Gender(male/female) | 64/60 | 47/46 | 17/14 | 0.83 |
| Tumor growth<br>pattern/N<br>(focal/diffuse) | 31/93 | 20/73 | 11/20 | 0.18 |
| Duration/months | 4.93(10.74) | 3.23(3.37) | 3.73(8.79) | 0.84 |
| KPS/scores | 79.49(14.69) | 74(17.29) | 77.79(14.87) | 0.83 |
| PFS/days | 558.91(534.07) | 451.22(368.63) | 487.96(313.17) | - |
| Tumor_volume/cm<sup>3</sup> | 27.44(14.58) | 30.76(15.97) | 27.01(12.68) | 0.83 |
| tOCP_CT/N | 4.17(4.10) | 7.23(5.25) | 3.81(3.71) | **0.000** |
| tOCN_CT/N | 3.96(4.86) | 1.08(1.07) | 3.77(3.81) | **0.000** |

Specifically, subtype 1 exhibited widespread positive deviations, primarily in the postcentral gyrus, temporal pole, middle frontal gyrus, and superior parietal lobule (Fig 5A, B), whereas subtype 2 demonstrated extensive negative deviations, mainly in the temporoparietal junction, inferior frontal gyrus, and inferior temporal gyrus (Fig 5A, C). Further intergroup analysis pinpointed the prefrontal lobe, superior frontal gyrus, superior temporal gyrus, middle temporal gyrus, supramarginal gyrus, and parietal lobe as regions of most significant difference between the two subtypes, particularly the precentral gyrus and parietal lobe (P_FDR_ < 0.05) (Fig 5D). These discoveries underscore the significant heterogeneity in cortical thickness deviation patterns among brainstem tumor subtypes.

**Fig 4.**
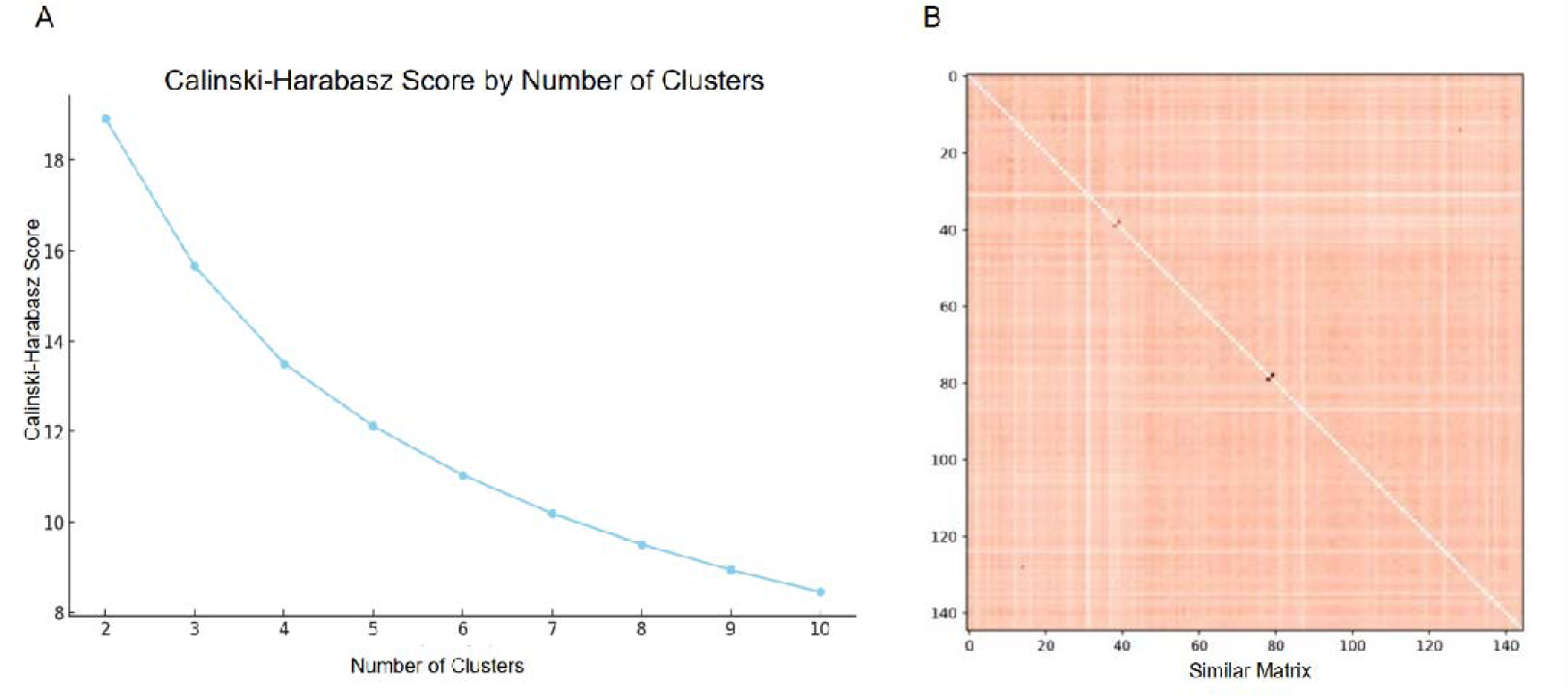
Evaluation index of spectral clustering. Calinski-Harabasz Score by Number of Clusters. This plot displays the Calinski-Harabasz score as a function of the number of clusters. The score decreases as the number of clusters increases from 2 to 10, indicating the quality of clustering for different numbers of clusters. (A) Calinski-Harabasz score for spectral clustering evaluation. (B) Similar matrix of spectral clustering. Similarity Matrix. The heatmap represents the similarity matrix used for clustering. The color intensity indicates the degree of similarity between different data points, with darker colors representing higher similarity. The diagonal represents perfect similarity, as each point is compared to itself.

**Fig 5.**
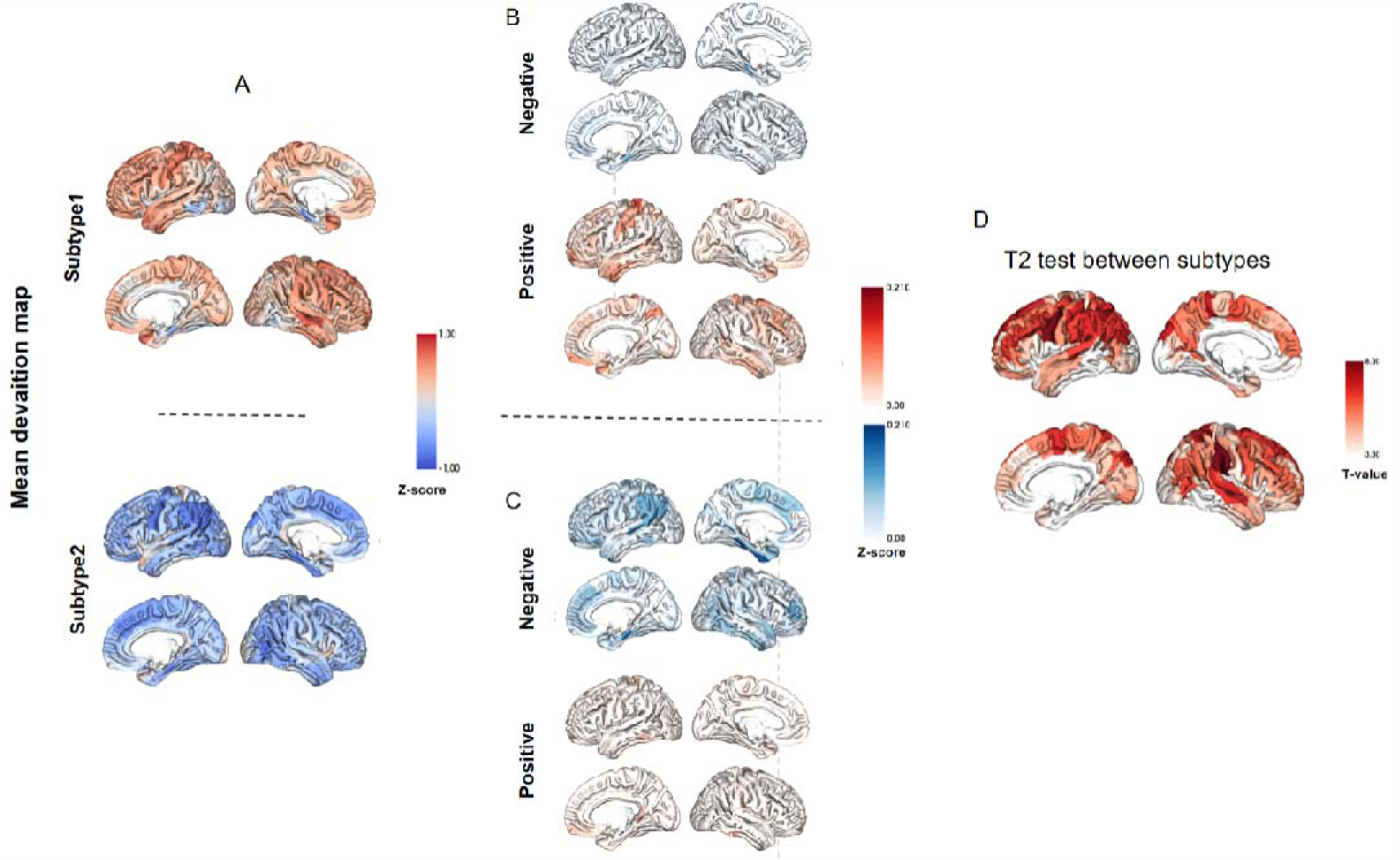
Spatial overlap map of deviations in subtypes. (A) The mean deviation map for two subtypes. (B) Positive and negative deviations in subtype 1. (C) Positive and negative deviations in subtype 2. (D) Differences between the two subtypes.

### Clinical Differences between Subtypes

Univariate analysis revealed no significant differences between the two brainstem tumor subtypes in terms of age, gender, tumor growth patterns, KPS, or tumor volume (see Table 2). However, the two-way ANOVA results revealed a significant main effect of subtype on duration (F_(1, 94)_ = 6.69, p = 0.011). Additionally, there was a significant main effect of tumor growth pattern on duration (F_(1, 94)_ = 4.36, p = 0.039). However, the interaction effect between subtype and DIPG on duration was not significant (F_(1, 94)_ = 2.80, p = 0.098) (Fig 6A). These results suggest that while both subtype and DIPG independently influence the duration, their interaction does not significantly affect it.

**Fig 6.**
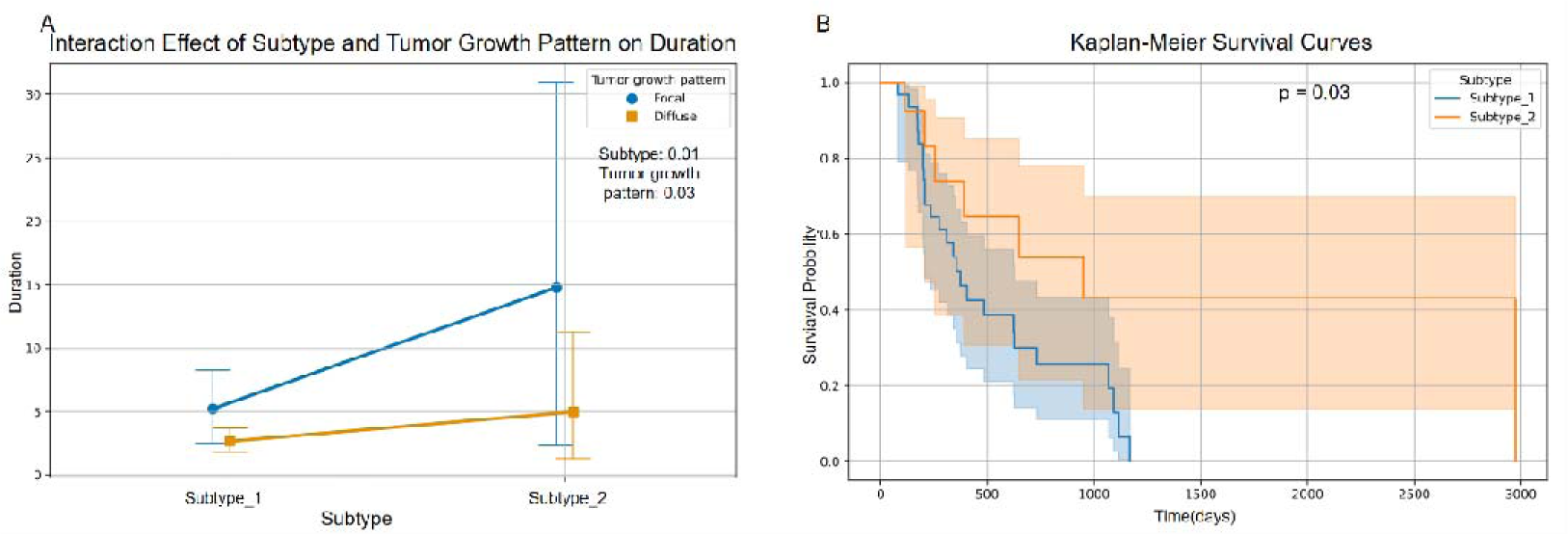
Clinical effects on subtypes. (A) Interaction effect of subtype and tumor growth pattern on duration. The plot shows the interaction between two subtypes (Subtype 1 and Subtype 2) and two tumor growth patterns (Focal and Diffuse) on the duration (in months). Subtype 1 with a focal growth pattern shows a significantly shorter duration compared to Subtype 2 with a focal growth pattern. The statistical significance is indicated for both the subtype (p = 0.01) and tumor growth pattern (p = 0.03). (B) Kaplan-Meier curves of subtypes. Comparing the survival probability of two subtypes (Subtype 1 and Subtype 2) over time (in days). The curves indicate a significant difference in survival between the two subtypes (p = 0.03), with Subtype 2 showing a higher survival probability over time. The shaded areas represent the 95% confidence intervals for each curve.

### Prognostic Factors and Survival Analysis

The Cox regression analysis of prognostic factors indicated that most variables were not statistically significant predictors of prognosis (see Table 3). Age, sex, tumor growth pattern, tOCP_CT, and tumor volume were all found to be non-significant predictors. Notably, tOCN_CT (coef = -0.13, 95% CI: 0.76-1.01, p = 0.06) approached significance, suggesting a potential trend where higher levels of tOCN_CT might be associated with a lower risk, though this did not reach conventional statistical significance. However, duration was a statistically significant predictor of prognosis (coef = -0.26, 95% CI: 0.62-0.95, p = 0.02), indicating that longer duration was associated with a lower risk of adverse outcomes.

**Table 3.**
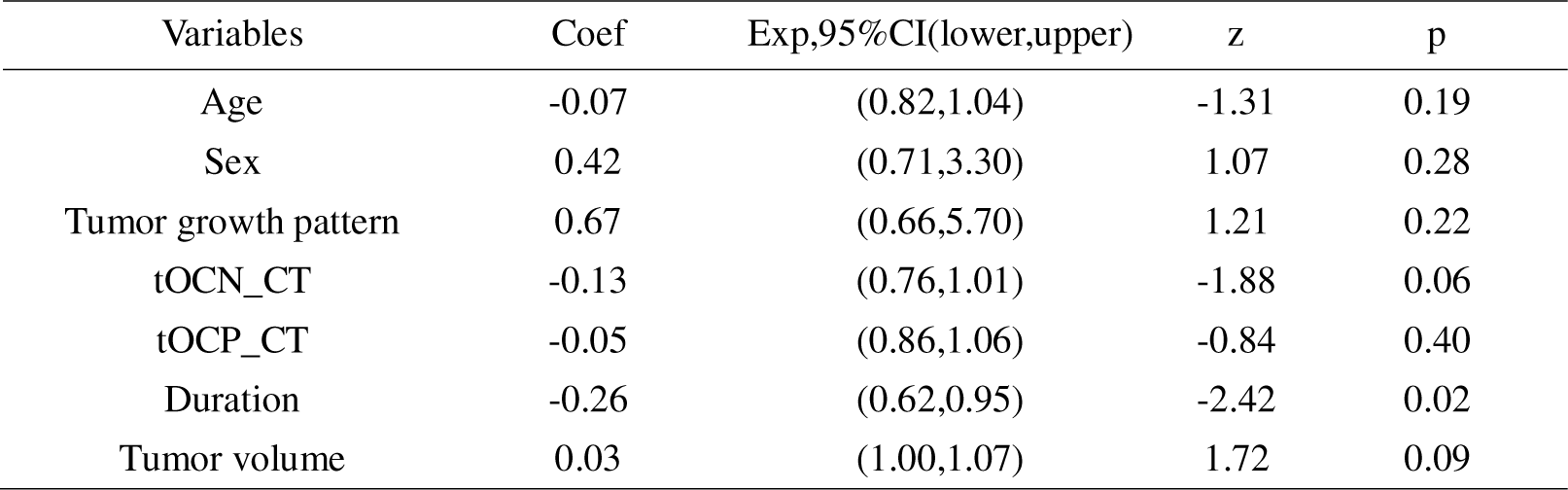
Cox regression of prognosis.

Furthermore, the Kaplan-Meier survival curves for subtype 1, subtype 2 and subtype 3 illustrate a significant difference in survival probabilities over time (p<0.05) (see Fig 6). Subtype 1 demonstrates a higher survival probability compared to subtype 2 and subtype 3 throughout the observed period. The shaded areas around the curves represent the 95% confidence intervals, indicating the reliability of the survival estimates for each subtype. This significant difference suggests that subtype 1 have a better prognosis.

## Discussion

Our study constructed normative growth models for cortical thickness, uncovering pronounced heterogeneous deviations in brainstem tumor patients. Specifically, focal tumors exhibited a correlation with cortical thinning, whereas diffuse tumors were associated with cortical thickening. Furthermore, spectral clustering analysis successfully classified patients into two distinct biological subtypes based on their unique cortical thickness deviation patterns. Notably, both subtype and tumor growth pattern exhibited a dependent significant influence on duration. Additionally, Subtype 2 demonstrated a higher survival probability compared to Subtype 1, indicating a better prognosis. These discoveries provide fresh perspectives on the diagnosis, treatment, and prognostic evaluation of brainstem tumors.

In this study, we observed significant heterogeneity in cortical thickness deviations across multiple brain regions in patients with brainstem tumors. This deviation was not only present between patients and healthy controls but also varied among individual patients. The heterogeneous cortical thickness deviations reflect the non-uniform impact of tumors on brain structure, potentially related to tumor growth patterns, locations, as well as individual genetic and physiological differences. Previous studies have explored morphological changes in distal brain regions induced by gliomas, such as increased gray matter volume in the contralateral insula in insular gliomas (Almairac et al, 2018) and cortical thickness reorganization in the superior parietal lobe of the contralateral hemisphere in frontal lobe gliomas (Liu et al, 2020).

Further exploring revealed a significant association between cortical thickness alterations and clinical features in brainstem tumor patients. Specifically, the correlation between focal tumors and cortical thinning, along with the positive correlation between tumor volume and cortical thickness deviations, suggests specific impacts of tumor growth patterns and sizes on cortical structure. This implies that cortical atrophy predominates in focal tumor patients, likely related to the focal growth characteristics of these tumors, which are visually confined to one side of the brain or manifest as exophytic tumors. Pathologically, these tumors encompass low-grade astrocytomas, hemangioblastomas, oligodendrogliomas, pilocytic astrocytomas, and ependymomas, sharing common features such as slow growth, primarily local proliferation rather than widespread infiltration, and relatively longer disease courses (D’aes & Mariën, 2014; Klimo et al, 2013). Notably, the average disease duration in focal tumor patients is significantly longer than that in diffuse tumor patients. Additionally, these focal tumo patients typically present with high intracranial pressure symptoms like headaches and nausea, with a minority exhibiting hydrocephalus, which is a crucial factor contributing to cortical thinning(Boes et al, 2015). Cortical thinning may be associated with multiple factors, including neuronal loss, synaptic pruning, and reactive gliosis (Altmann et al, 2021), underlying the pathophysiological changes involved.

On the other hand, cortical changes in diffuse tumor patients were primarily characterized by thickening, consistent with previous findings in patients with diffuse midline glioma (DMG). DMG patients exhibit widespread damage to myelinated axons in the cerebral cortex, and these patients show thicker cortical thickness in regions of the sensorimotor, salience, and default mode networks(Zhang et al, 2024). The cortical thickening observed in diffuse tumor patients may be attributed to the infiltrative ability of high-grade glioma cells into myelinated axons, leading to compensatory cortical thickening due to myelin loss(Douw et al). Furthermore, glioma cells may be distributed throughout the brain, and their invasion of brain regions can trigger disinhibition of nearby neural networks (Sahm et al, 2012; Yang et al, 2022). Structural plasticity can induce changes in dendrites, axons, and myelin sheaths, resulting in cortical thickening. This cortical thickening may stem from various factors, including larger cells, higher columnar structure density, the generation of new neuroglial cells, or a series of changes such as myelin plasticity, axonal sprouting, or angiogenesis (Xin & Chan, 2020). Additionally, it is well-known that the integrity of white matter fibers is disrupted in glioma patients, and white matter diffusion metrics are correlated with early infiltration in diffuse gliomas (Jutten et al, 2019). Not only is the white matter microstructure skeleton in the vicinity of the tumor damaged, but white matter destruction can also be observed in the contralateral hemisphere, suggesting that chronic damage caused by gliomas may lead to the degeneration of distal fiber tracts.(Zhang et al, 2023)These findings are crucial for assessing disease severity, predicting treatment responses, and prognoses. For instance, patients with significant cortical thinning may face higher risks of neurological impairment, while those with larger tumors may require more aggressive interventions. Thus, incorporating cortical thickness changes into clinical assessments can enhance our comprehensive understanding of patient disease states and facilitate more precise treatment planning.

A core finding of this study is the significant variability in cortical thickness deviations among brainstem tumor patients within subtype classifications, highlighting the crucial value of cortical thickness as a potential imaging biomarker. Previous research has predominantly focused on the local characteristics of tumors to predict survival, often relying on segmentation algorithms that necessitate large training datasets(Feng et al, 2020; Sun et al, 2019). Despite continuous exploration of new imaging markers, such as tumor-induced deformation of surrounding brain structures and connectome(Wei et al, 2023; Wittenberg, 2010), the need for innovative predictive indicators persists. Cortical thickness has been demonstrated to be associated with the diagnosis and prognosis of neurological and psychiatric disorders. However, its application in routine clinical practice has been limited due to the lack of normative comparisons(Nagtegaal et al, 2020; Tahedl, 2020). This study uniquely underscores the potential value of cortical thickness changes in survival prediction from a whole-brain structural perspective. The significant difference in survival between the subtypes underscores the importance of considering cortical thickness deviations as potential biomarkers for prognosis in brainstem tumor patients. These findings highlight the need for further research to elucidate the biological underpinnings of these associations and to explore the potential of cortical thickness measurements in guiding personalized treatment strategies. Subtype classification not only deepens our understanding of the heterogeneity of brainstem tumors but also provides a new basis for precise diagnosis of the disease. As a crucial indicator of brain structure, changes in cortical thickness reflect both the direct and indirect impacts of tumors on brain tissue. Therefore, incorporating cortical thickness deviations into the imaging assessment system can enhance our comprehension of the pathophysiological processes of brainstem tumors and offer robust support for the development of personalized treatment plans.

This study enhances our understanding of the heterogeneity in brainstem tumor patients and underscores the prognostic value of cortical thickness deviations. It holds significant implications for personalized treatment and clinical practice in brainstem tumors. Subtype classification allows for tailored therapeutic interventions, potentially improving treatment efficacy and reducing side effects. It also aids in accurate prognosis prediction, enabling early identification of high-risk patients for preventive measures and closer monitoring. Additionally, it offers a novel perspective for brainstem tumor research, facilitating deeper insights into tumor pathogenesis and progression.

However, the study has limitations that need to be addressed in future research. The findings should be validated in larger cohorts to ensure generalizability. Additionally, the study primarily focuses on cortical thickness deviations, and other factors such as genetic and molecular characteristics should be considered to fully understand the underlying mechanisms driving these associations. Further research should also explore the integration of cortical thickness measurements with other imaging biomarkers to enhance prognostic accuracy and treatment strategies.

## Conclusions

In conclusion, this study reveals two major patterns of cortical thickness changes induced by the heterogeneity of brainstem tumors. This finding deepens our understanding of the relationship between brainstem tumors and cortical morphology. It also elucidates the complex interactions between tumor size, growth patterns, and cortical thickness alterations. Importantly, the results suggest that changes in cortical thickness could serve as valuable biomarkers for predicting progression-free survival, which is significant for clinical assessment and treatment strategy formulation. Therefore, this study provides new perspectives and potential targets for personalized treatment of brainstem tumor patients. It also sheds light on the physiological mechanisms by which brainstem tumors affect brain structure, laying a theoretical foundation for more precise clinical interventions and efficacy monitoring in the future.

## Author Contributions

Liwei Zhang and Lingzhong Fan designed the task. Heyuan Jia and Kaikai Wang analyzed all data and edited the manuscript. Peng Zhang revised the manuscript. Congying Chu checked the research method. Mingxin Zhang and Yiying Mai helped collected the data. Xuntao Yin helped collected data of healthy children.

## Reference

Acharya, A., Ren, P., Yi, L., Tian, W. & Liang, X. (2022) Structural atrophy and functional dysconnectivity patterns in the cerebellum relate to cerebral networks in svMCI. Front Neurosci, 16, 1006231.

Almairac, F., Duffau, H. & Herbet, G. (2018) Contralesional macrostructural plasticity of the insular cortex in patients with glioma: A VBM study. Neurology, 91(20), e1902–e1908.

Altmann, A., Ryten, M., Di Nunzio, M., Ravizza, T., Tolomeo, D., Reynolds, R. H., Somani, A., Bacigaluppi, M., Iori, V., Micotti, E., Di Sapia, R., Cerovic, M., Palma, E., Ruffolo, G., Botía, J. A., Absil, J., Alhusaini, S., Alvim, M. K. M., Auvinen, P., Bargallo, N., Bartolini, E., Bender, B., Bergo, F. P. G., Bernardes, T., Bernasconi, A., Bernasconi, N., Bernhardt, B. C., Blackmon, K., Braga, B., Caligiuri, M. E., Calvo, A., Carlson, C., Carr, S. J. A., Cavalleri, G. L., Cendes, F., Chen, J., Chen, S., Cherubini, A., Concha, L., David, P., Delanty, N., Depondt, C., Devinsky, O., Doherty, C. P., Domin, M., Focke, N. K., Foley, S., Franca, W., Gambardella, A., Guerrini, R., Hamandi, K., Hibar, D. P., Isaev, D., Jackson, G. D., Jahanshad, N., Kälviäinen, R., Keller, S. S., Kochunov, P., Kotikalapudi, R., Kowalczyk, M. A., Kuzniecky, R., Kwan, P., Labate, A., Langner, S., Lenge, M., Liu, M., Martin, P., Mascalchi, M., Meletti, S., MoritalJSherman, M. E., O’Brien, T. J., Pariente, J. C., Richardson, M. P., RodriguezlJCruces, R., Rummel, C., Saavalainen, T., Semmelroch, M. K., Severino, M., Striano, P., Thesen, T., Thomas, R. H., Tondelli, M., Tortora, D., Vaudano, A. E., Vivash, L., von Podewils, F., Wagner, J., Weber, B., Wiest, R., Yasuda, C. L., Zhang, G., Zhang, J., Leu, C., Avbersek, A., Thom, M., Whelan, C. D., Thompson, P., McDonald, C. R., Vezzani, A. & Sisodiya, S. M. (2021) A systemslJlevel analysis highlights microglial activation as a modifying factor in common epilepsies. Neuropathology and Applied Neurobiology, 48(1).

Boes, A. D., Prasad, S., Liu, H., Liu, Q., Pascual-Leone, A., Caviness, V. S., Jr. & Fox, M. D. (2015) Network localization of neurological symptoms from focal brain lesions. Brain, 138(Pt 10), 3061–75.

Boukaka, R. G., Beuriat, P.-A., Di Rocco, F., Vasiljevic, A., Szathmari, A. & Mottolese, C. (2023) Brainstem tumors in children: a monocentric series in the light of genetic and bio-molecular progress in pediatric neuro-oncology. Frontiers in Pediatrics, 11.

Bu, X., Zhao, Y., Zheng, X., Fu, Z., Zhang, K., Sun, X., Cui, Z., Xia, M., Ma, L., Liu, N., Lu, J., Zhao, G., Ding, Y., Deng, Y., Wang, J., Chen, R., Zhang, H., Men, W., Wang, Y., Gao, J., Tan, S., Sun, L., Qin, S., Tao, S., Wang, Y., Dong, Q., Cao, Q., Yang, L. & He, Y. (2024) Normative growth modeling of brain morphology reveals neuroanatomical heterogeneity and biological subtypes in children with ADHD.

Chieffo, D. P. R., Lino, F., Ferrarese, D., Belella, D., Della Pepa, G. M. & Doglietto, F. (2023) Brain Tumor at Diagnosis: From Cognition and Behavior to Quality of Life. Diagnostics (Basel), 13(3).

Chou, S. C., Chen, Y. N., Huang, H. Y., Kuo, M. F., Wong, T. T., Kuo, S. H. & Yang, S. H. (2024) Contemporary Management of Pediatric Brainstem Tumors. Adv Tech Stand Neurosurg, 49, 231–254.

D’aes, T. & Mariën, P. (2014) Cognitive and Affective Disturbances Following Focal Brainstem Lesions: A Review and Report of Three Cases. The Cerebellum, 14(3), 317–340.

Douw, L. A.-O., Breedt, L. C. & Zimmermann, M. L. M. Cancer meets neuroscience: the association between glioma occurrence and intrinsic brain features(1460-2156 (Electronic)).

Duffau, H. (2014) Diffuse low-grade gliomas and neuroplasticity. Diagn Interv Imaging, 95(10), 945–55.

Fan, L., Li, H., Zhuo, J., Zhang, Y., Wang, J., Chen, L., Yang, Z., Chu, C., Xie, S., Laird, A. R., Fox, P. T., Eickhoff, S. B., Yu, C. & Jiang, T. (2016) The Human Brainnetome Atlas: A New Brain Atlas Based on Connectional Architecture. Cereb Cortex, 26(8), 3508–26.

Feng, X., Tustison, N. J., Patel, S. H. & Meyer, C. H. (2020) Brain Tumor Segmentation Using an Ensemble of 3D U-Nets and Overall Survival Prediction Using Radiomic Features. Front Comput Neurosci, 14, 25.

Fortin, J. P., Cullen, N., Sheline, Y. I., Taylor, W. D., Aselcioglu, I., Cook, P. A., Adams, P., Cooper, C., Fava, M., McGrath, P. J., McInnis, M., Phillips, M. L., Trivedi, M. H., Weissman, M. M. & Shinohara, R. T. (2018) Harmonization of cortical thickness measurements across scanners and sites. Neuroimage, 167, 104–120.

Jutten, K., Mainz, V., Gauggel, S., Patel, H. J., Binkofski, F., Wiesmann, M., Clusmann, H. & Na, C. H. (2019) Diffusion Tensor Imaging Reveals Microstructural Heterogeneity of Normal-Appearing White Matter and Related Cognitive Dysfunction in Glioma Patients. Front Oncol, 9, 536.

Keller, S. S. & Roberts, N. (2008) Voxel-based morphometry of temporal lobe epilepsy: an introduction and review of the literature. Epilepsia, 49(5), 741–57.

Klimo, P., Jr., Pai Panandiker, A. S., Thompson, C. J., Boop, F. A., Qaddoumi, I., Gajjar, A., Armstrong, G. T., Ellison, D. W., Kun, L. E., Ogg, R. J. & Sanford, R. A. (2013) Management and outcome of focal low-grade brainstem tumors in pediatric patients: the St. Jude experience. J Neurosurg Pediatr, 11(3), 274–81.

Lee, B. C., Kneeland, J. B., Walker, R. W., Posner, J. B., Cahill, P. T. & Deck, M. D. (1985) MR imaging of brainstem tumors. AJNR Am J Neuroradiol, 6(2), 159–63.

Liu, Y., Hu, G., Yu, Y., Jiang, Z., Yang, K., Hu, X., Li, Z., Liu, D., Zou, Y., Liu, H. & Chen, J. (2020) Structural and Functional Reorganization Within Cognitive Control Network Associated With Protection of Executive Function in Patients With Unilateral Frontal Gliomas. Front Oncol, 10, 794.

Marquand, A. F., Rezek, I., Buitelaar, J. & Beckmann, C. F. (2016) Understanding Heterogeneity in Clinical Cohorts Using Normative Models: Beyond Case-Control Studies. Biol Psychiatry, 80(7), 552–61.

Nagtegaal, S. H. J., David, S., Snijders, T. J., Philippens, M. E. P., Leemans, A. & Verhoeff, J. J. C. (2020) Effect of radiation therapy on cerebral cortical thickness in glioma patients: Treatment-induced thinning of the healthy cortex. Neurooncol Adv, 2(1), vdaa060.

Newman, D. B. (1995) Anatomy and neurotransmitters of brainstem motor systems. Adv Neurol, 67, 219–44.

Novak, V., Novak, P., deMarchie, M. & Schondorf, R. (1995) The effect of severe brainstem injury on heart rate and blood pressure oscillations. Clin Auton Res, 5(1), 24–30.

Osswald, M., Jung, E., Sahm, F., Solecki, G., Venkataramani, V., Blaes, J., Weil, S., Horstmann, H., Wiestler, B., Syed, M., Huang, L., Ratliff, M., Karimian Jazi, K., Kurz, F. T., Schmenger, T., Lemke, D., Gommel, M., Pauli, M., Liao, Y., Haring, P., Pusch, S., Herl, V., Steinhauser, C., Krunic, D., Jarahian, M., Miletic, H., Berghoff, A. S., Griesbeck, O., Kalamakis, G., Garaschuk, O., Preusser, M., Weiss, S., Liu, H., Heiland, S., Platten, M., Huber, P. E., Kuner, T., von Deimling, A., Wick, W. & Winkler, F. (2015) Brain tumour cells interconnect to a functional and resistant network. Nature, 528(7580), 93–8.

Pistoia, F., Cornia, R., Conson, M., Gosseries, O., Carolei, A., Sacco, S., Quattrocchi, C. C., Mallio, C. A., Iani, C., Mambro, D. D. & Sara, M. (2016) Disembodied Mind: Cortical Changes Following Brainstem Injury in Patients with Locked-in Syndrome. Open Neuroimag J, 10, 32–40.

Recinos, P. F., Sciubba, D. M. & Jallo, G. I. (2007) Brainstem tumors: where are we today? Pediatr Neurosurg, 43(3), 192–201.

Sahm, F., Capper, D., Jeibmann, A., Habel, A., Paulus, W., Troost, D. & von Deimling, A. (2012) Addressing Diffuse Glioma as a Systemic Brain Disease With Single-Cell Analysis. Archives of Neurology, 69(4), 523–526.

Smith, L. H. & DeMyer, W. E. (2003) Anatomy of the brainstem. Semin Pediatr Neurol, 10(4), 235–40.

Sun, L., Zhang, S., Chen, H. & Luo, L. (2019) Brain Tumor Segmentation and Survival Prediction Using Multimodal MRI Scans With Deep Learning. Front Neurosci, 13, 810.

Tahedl, M. (2020) Towards individualized cortical thickness assessment for clinical routine. J Transl Med, 18(1), 151.

Wang, X., Zhou, C., Wang, Y. & Wang, L. (2022) Microstructural changes of white matter fiber tracts induced by insular glioma revealed by tract-based spatial statistics and automatic fiber quantification. Sci Rep, 12(1), 2685.

Wei, Y., Li, C., Cui, Z., Mayrand, R. C., Zou, J., Wong, A., Sinha, R., Matys, T., Schonlieb, C. B. & Price, S. J. (2023) Structural connectome quantifies tumour invasion and predicts survival in glioblastoma patients. Brain, 146(4), 1714–1727.

Wittenberg, G. F. (2010) Experience, cortical remapping, and recovery in brain disease. Neurobiol Dis, 37(2), 252–8.

Xi-Nian, Z. & Consortium, C. (2024) developing Chinese Color Nest Project (devCCNP) Lite. 01/19.

Xin, W. & Chan, J. R. (2020) Myelin plasticity: sculpting circuits in learning and memory. Nat Rev Neurosci, 21(12), 682–694.

Yang, Y., Schubert, M. C., Kuner, T., Wick, W., Winkler, F. & Venkataramani, V. (2022) Brain Tumor Networks in Diffuse Glioma. Neurotherapeutics, 19(6), 1832–1843.

Zhang, S., Yang, X., Tan, Q., Sun, H., Chen, D., Chen, Y., Zhang, H., Yang, Y., Gong, Q. & Yue, Q. (2024) Cortical myelin and thickness mapping provide insights into whole-brain tumor burden in diffuse midline glioma. Cereb Cortex, 34(1).

Zhang, S., Zhao, F., Yang, X., Tan, Q., Li, S., Shao, H., Su, X., Gong, Q. & Yue, Q. (2023) Multiparametric mapping of white matter reorganizations in patients with frontal glioma-related epilepsy. CNS Neurosci Ther, 29(8), 2366–2376.

